# Salmonella exploits the reactive oxygen species generated by the plant immune system to enhance colonization

**DOI:** 10.1101/2024.03.04.583411

**Authors:** M Srai Aldin, E Tzipilevich

## Abstract

Understanding bacterial interaction with the plant immune system is essential for building a sustainable food system. Discouraging pathogens enhancing beneficial bacteria and preventing food spoilage and contamination by human enteric pathogens. Immune system activation triggers a burst of toxic reactive oxygen species (ROS). We discovered that the food-poisoning bacterium *Salmonella* exploits the plant ROS to enhance biofilm formation and colonization. Scavenging the ROS with antioxidants or inhibiting their production reduces colonization. Further, combining the antioxidant vitamin C and immune system activation with salicylic acid synergistically inhibited *Salmonella* colonization of the plant model Arabidopsis and edible plants. This safe-to-use combination is an eco-friendly method to prevent plant contamination by *Salmonella, Listeria* and enteropathogenic *E. coli*. Finally, we have found that exploiting the plant immunity for enhancing colonization is not exclusive to *Salmonella* and diverse beneficial *pseudomonas* species isolated from plants can also exploit the plant immune system, suggesting that it is a general plant colonization strategy.

## Introduction

The increase in the world population, global warming and the reduction in the efficiency of chemical fertilizers and pesticides necessitate the building of sustainable food systems^1^. Understanding plant microbe interaction at the molecular level is an avenue to reach this goal. Better understanding will enable us to discourage pathogens, enhance beneficial bacteria and prevent food spoilage and contamination by human enteric pathogens^2^.

Food poisoning by enteric pathogens is a leading cause of morbidity and mortality worldwide^3^. The increased consumption of fresh fruits and vegetables (fresh produce), due to awareness of their importance to a healthy lifestyle and the extension of the supply chains, made it a significant source of food-borne illnesses. The Gram-negative bacteria *Salmonella* and *E. coli* and the Gram-positive bacterium *Listeria monocytogenes* are the three main bacteria contaminating fresh produce^3-5^. Chemical disinfection of fruits and vegetables adversely affects workers’ and consumers’ health. Thus, eco-friendly methods to ensure food safety are highly needed^6^.

Research from recent years revealed that human pathogens can colonize plants and may even utilize plants as a secondary host^5,7^. They exhibit a complex interaction with plants and activate plant immune responses. Thus, understanding their plant colonization process could inform rational approaches to fight food contamination. In this regard, utilizing plant immunity to fight salmonella contamination is extensively explored^6,8^. We have analyzed the molecular interaction between the enteropathogenic bacteria *Salmonella enterica* and the plant model Arabidopsis thaliana. As previously shown, the *Salmonella* activates the plant immune system^9-11^, despite exhibition of beneficial lifestyle on Arabidopsis with no obvious pathogenic symptoms. The first level of plant immunity comprises receptors that recognize conserved microbial molecules, e.g., flagella, elongation factor Tu, prevalent in both plant pathogens and non-pathogenic bacteria. Receptor activation leads to a burst of reactive oxygen species (ROS) and a signal transduction cascade^12^. Specialized plant pathogens employ a plethora of effectors that target plant immunity, mainly delivered through the type III secretion system. Thus, they can evade the first protection line and grow to a high number^13^. However, if and how ‘non-specialized’ plant pathogens including commensal and beneficial bacteria evade immune system toxicity is much less understood^14^.

Bacteria synthesize extracellular matrix to facilitate surface adhesion^15^. The matrix comprises sugar polymers, protein fibres, extracellular DNA and cell debris. Bacteria encased in a matrix are known as biofilm^16^. The matrix protects the bacteria from chemical assaults and, as such, poses a serious challenge for antibiotic treatment^17^. Biofilm formation is essential for root colonization^15,18^. However, the effect of the biofilm matrix on bacteria dealing with the plant immune system is not explored in detail. Our research revealed that biofilm formation protects *Salmonella* from the reactive oxygen species (ROS) produced upon plant immune system activation. Further, the bacteria exploited the ROS to enhance biofilm formation and colonization. Thus, immune system recognition paradoxically enhances root colonization.

Revealing the importance of ROS for facilitating biofilm formation and colonization enabled us to outsmart the bacteria. Scavenging the ROS with antioxidants reduced bacterial colonization and sensitized the bacteria to the plant immune system activity. The combination of the antioxidant ascorbic acid (vitamin C) and excessive immune system activation by the plant hormone salicylic acid (SA) exhibited synergistic inhibition of bacterial contamination. Further, the combination of vitamin C and SA inhibits plant colonization of various *salmonella* serovars, *listeria monocytogenes* and enteropathogenic *E. coli*, both in Arabidopsis and in edible plants, making it a promising method to ensure food safety. Finally, we bring initial evidence that exploitation of the plant immunity for enhanced colonization is utilized by other bacteria from the pseudomonas genus.

## Results

### *Salmonella* exhibited reduced colonization of immune mutant plants

We inoculated seedlings with 10^6^ CFU/ml of *Salmonella* (the concentration of inoculated bacteria from here on unless otherwise stated), we used *Salmonella enterica Serovar Typhimurium 14028s* throughout the paper except for specific experiments. Notably, at this number the bacteria exhibited beneficial effect and significantly enhanced primary root length, while 10^8^ CFU/ml inhibited root length (Figure S1A), consistent with previous results showing plant pathogenic effect of *Salmonella* at high concentration [e.g.^11^]. Despite the beneficial effect *Salmonella* stimulated immune system activation in Arabidopsis (Figure 1A and Figure S1B), as judged by FrK1 promoter expression, a downstream target of the FLS2 receptor^19^ (Figure 1A) and CYP71A12 promoter expression, a gene involve in biosynthesis of the secondary metabolite camalexin^20^ (Figure S1B). To understand how the plant immune system modulates bacterial colonization, we inoculated bacteria into bbc (*bak1,bkk1,cerk1*) triple mutant plants, lacking all three co-receptors of the pattern-triggered immunity^21^. We anticipated overgrowth of *Salmonella* on plants perturbed in immune recognition. However, when inoculated with 10^6^ CFU/ml bacteria exhibited no significant reduction [12% reduction on bbc compared to Col0 but P = 0.07) (Figure 1B)]. Immune system activation triggers a burst of ROS production to kill the invading pathogen^22,23^. However, inoculating *Salmonella* into *rbohd rbohf* plants perturbed in ROS production in response to immune system activation^24^ revealed a significant reduction rather than an increase in root colonization (Figure 1B), [compare for example to^23^ where Bacilli overgrown on *rbohd rbohf* plants]. Of note, when seedlings inoculated with 10^8^ CFU bacteria exhibited significant overgrowth on bbc mutant plants (Figure S1C). It has been previously shown that *Salmonella* colonizes the endophyte of Arabidopsis^11,25^. However, we failed to detect any CFU from crashed roots after surface-sterilization (we performed 2 experiments with 4 roots in each with zero bacteria detected in all the roots).

**Figure 1.**
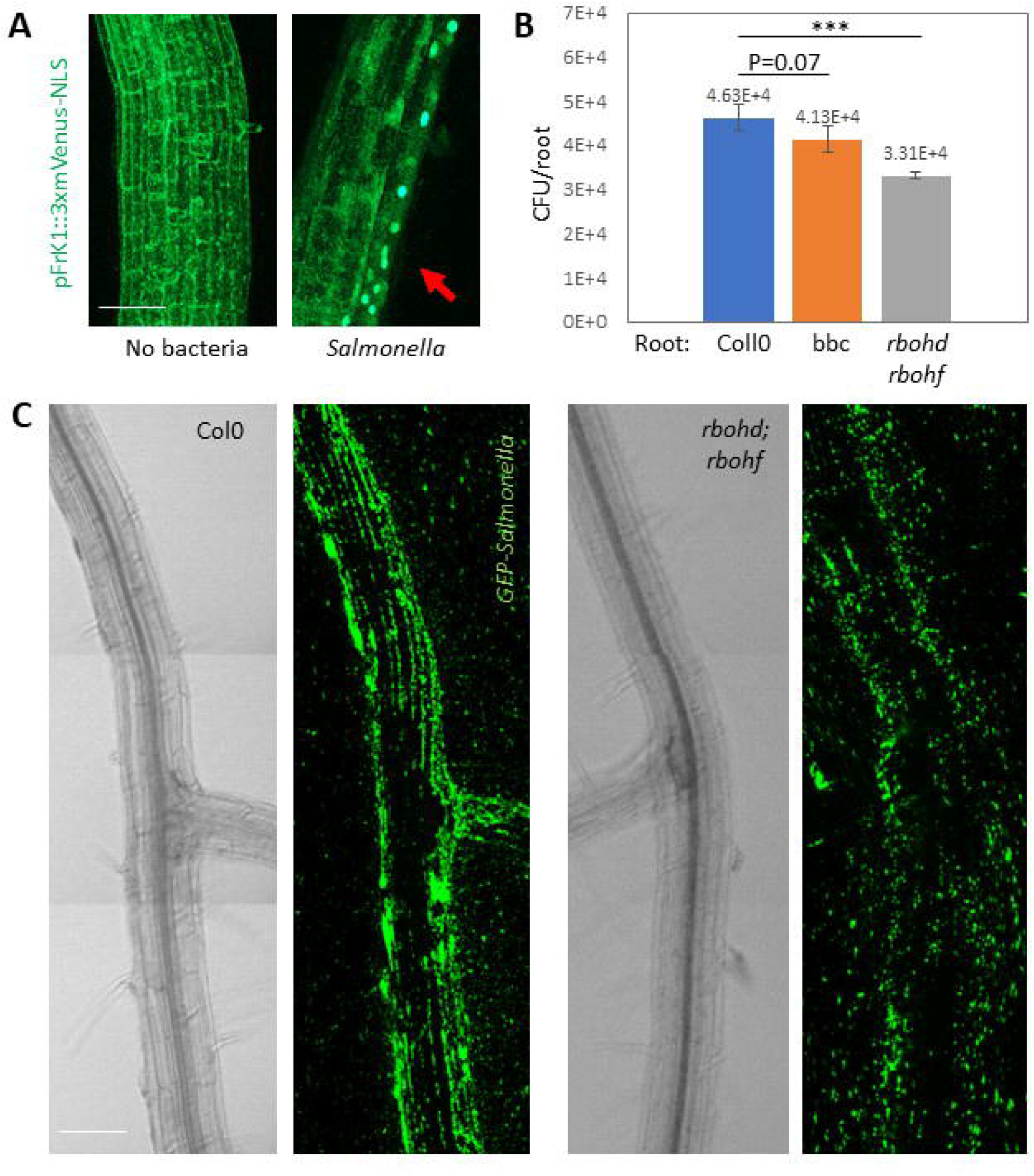
Immune system activation enhances root colonization by *Salmonella*. **(A)** days old Arabidopsis seedlings (pFrk1::3xmVenus-NLS) were inoculated with bacteria on MS agar plates. mVenus fluorescence was monitored after 18 hrs using confocal microscopy. Shown are representative maximal projection x20 fluorescent images out of 4 roots from each treatment. Red arrow mark nuclear localized mVenus expression, scale bar = 25μm. **(B)** Seedlings of the indicated genotypes were inoculated with *Salmonella*, and the number of colonizing bacteria was determined after 48 hrs. Shown are averages and standard deviations (SD) of bacterial colonization from at least 2 independent experiments with 4 seedlings in each. *** = P < 0.005. **(C)** *Salmonella* constitutively expressing GFP were inoculated into seedlings of the indicated genotypes for 48 hrs. Root colonization was monitored by confocal microscopy. Shown are representative maximal projection x20 fluorescent images and the complementary brightfield images of the same roots, out of 4 roots from each treatment. Bacteria colonizing WT roots (Col0) appeared in long stripes, while on roots mutant in ROS production (*rbohd;rbohf*) they did not form these strips, scale bar = 50μm.

To understand why bacteria failed to properly colonize *rbohd rbohf* plants, we monitored root colonization by GFP-expressing bacteria. On WT roots, bacteria appeared in long strips (Figure 1C). However, on *rbohd rbohf* plants, bacteria failed to form strips (Figure 1C). Thus, our results suggest that at low number of bacteria *Salmonella* can resist the activity of the plant immune system and that ROS production by the plant can even enhance the bacterial colonization (we discuss the differences between low and high number in the Discussion section).

### Biofilm formation is stimulated by the plant immune system derived ROS and protects the bacteria from the ROS toxicity

Biofilm formation is crucial for bacteria to adhere to plants^18^. We examined the colonization of three biofilm mutant strains: 1. *ΔcsgD*, a master regulator of curli polymers^26^2. *ΔbcsA*, encoding a cellulose synthase and 3. *ΔcsgB* a minor curli subunit plus *ΔbcsA* double mutant cells^*26-28*^. We found that mutant cells exhibited a significant reduction in root colonization (Figure 2A and Figure S2A-S2B), while PcsgD cells constitutively expressing CsgD^26^ exhibited a significant increase (Figure 2A), confirming the importance of biofilm formation for root colonization. Notably, the inability of *Salmonella* to form large stripes and patches on *rbohd rbohf* plants was reminiscent of bacteria perturbed in biofilm formation (Figure S2B). We hypothesized that plant ROS generated by RbohD and RbohF stimulate biofilm formation on the root. This idea is consistent with the ability of ROS to enhance biofilm in vitro (Figure 2B). Consistently, the colonization of *ΔcsgD* cells isn’t reduced when colonizing *rbohd rbohf* plants (Figure 2C). On the contrary, *ΔcsgD* cells exhibited a significant increase in the colonization of *rbohd rbohf* and bbc plants (Figure 2C), as well as *ΔbcsA* and *ΔbcsA ΔcsgB* cells (Figure S2A). Further, treatment of the roots with Diphenyliodonium (DPI), an inhibitor of the NADPH oxidase enzymes rbohD and rbohF^23^ reduced the growth of WT cells while increase the growth of *ΔcsgD* cells (Figure S2C-S2D). These results indicate that the immune system can inhibit the growth of *Salmonella*. However, the ability to form biofilm protected the WT bacteria from the ROS.

**Figure 2.**
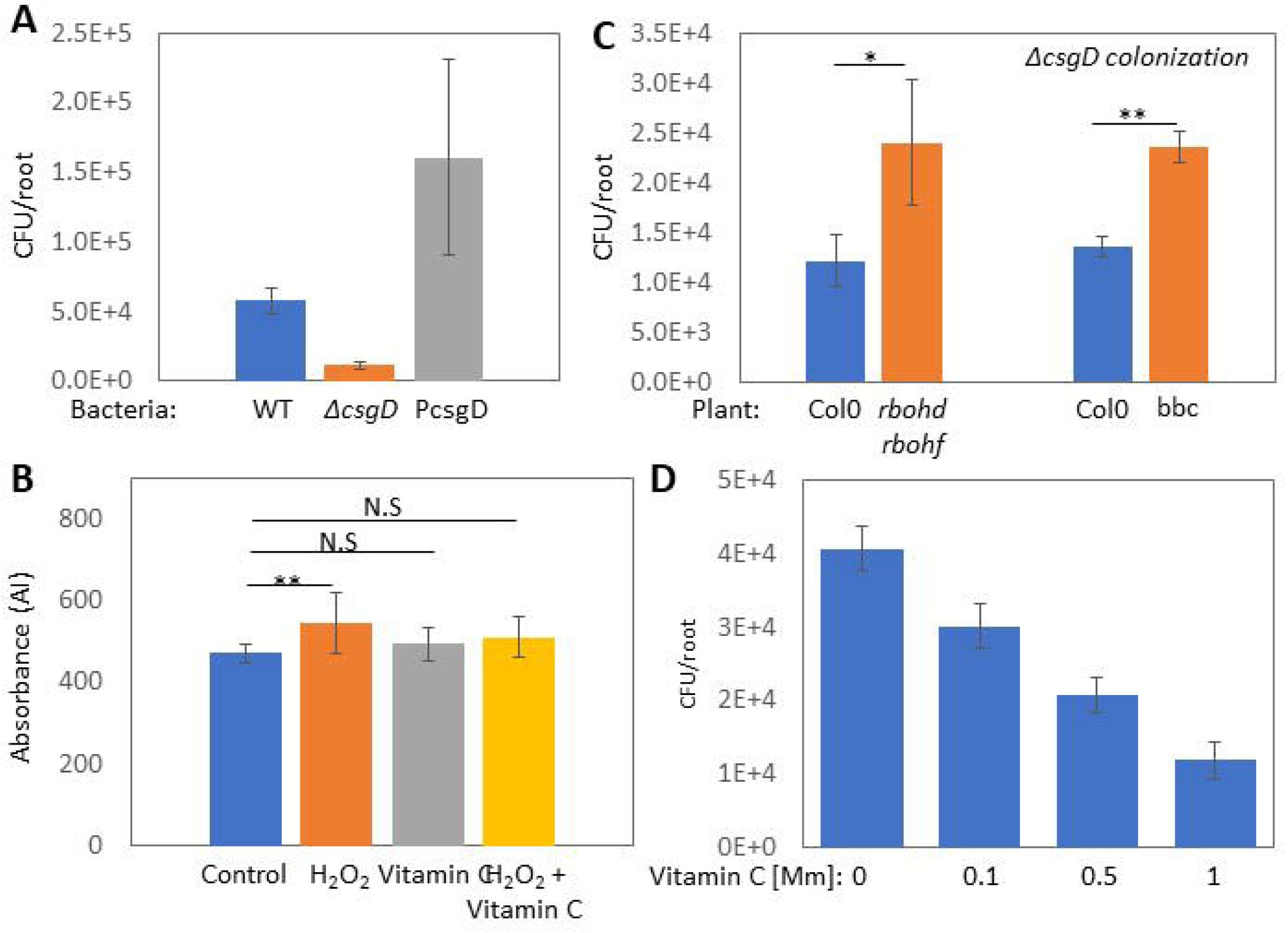
ROS enhances biofilm formation on the root. **(A)** 7 days old seedlings were inoculated with the indicated *Salmonella* strains on MS agar plates, and the number of colonizing bacteria was determined after 48 hrs. Shown are averages and standard deviations (SD) of bacterial colonization from 3 seedlings. **(B)** Bacteria inoculated into LB with no NaCl with the addition of 100μM H_2_O_2_, 1mM vitamin C, a combination of both or no addition, in a multi-well plate. The plate was kept with no shaking at 23° for 5 days, and adhesion to the well was monitored by crystal violet staining as an indicator of biofilm matrix formation. Shown are average and SD of at least 6 wells, * = P < 0.05. **(C)** Seedling of WT and the indicated immune mutant plants were inoculated with ΔcsgD bacteria, and the number of colonizing bacteria was determined after 48 hrs. Shown are averages and standard deviations (SD) of bacterial colonization from at least 2 independent experiments with 4 seedlings in each. * = P < 0.05, ** = P <0.01. ΔcsgD bacteria exhibited a significant increase in colonization on immune system mutants, in contrast to the decrease exhibited by WT bacteria. **(D)** Seedlings were inoculated with bacteria on control plates or plates with the indicated of vitamin C and the number of colonizing bacteria was determined after 48 hrs. Shown are averages and standard deviations (SD) of bacterial colonization from at least 2 experiments with 4 seedlings in each. All the treatments lead to a significant reduction in root colonization P < 0.05.

The results so far suggested that ROS positively affect salmonella colonization. Thus, we wondered if chemical ROS scavengers will inhibit colonization. Indeed, vitamin C and potassium iodide (KI) inhibited *Salmonella* colonization in a concentration dependent manner (Figure 2D and figure S3A-S3B). We further concentrated on vitamin C, which is a common antioxidant and is considered safe for human consumption even at high doses. Of note, the effect of vitamin C was independent of its acidity as we titrated the pH back to normal. Vitamin C did not affect biofilm formation *in vitro* (Figure 2B), but it can at least partially abolish the increase observed upon H_2_O_2_ addition (Figure 2B).

### Salicylic acid and vitamin C synergistically inhibited *Salmonella* colonization of Arabidopsis and edible plants

The results so far indicate that *Salmonella* exploits plant immunity to enhance biofilm formation. Nevertheless, excessive stimulation of the plant immunity using the stress hormone salicylic acid (SA) significantly inhibited *salmonella* colonization (Figure 3A-3B), [see also^6^]. Of note, SA did not affect the colonization of pCsgD, suggesting that robust biofilm matrix can protect the cells even from SA activity (Figure 3B). We next examined the application of vitamin C and SA together. Applying both chemicals together (50μM SA and 1Mm vitamin C) exhibited synergistic inhibition of salmonella colonization (Figure 3C) (The observed reduction is significantly lower than the theoretical combination of vitamin C x SA, P<0.05). Neither the compounds nor their combination affects bacterial growth in vitro (Figure 3D), indicating that their effect is plant specific.

**Figure 3.**
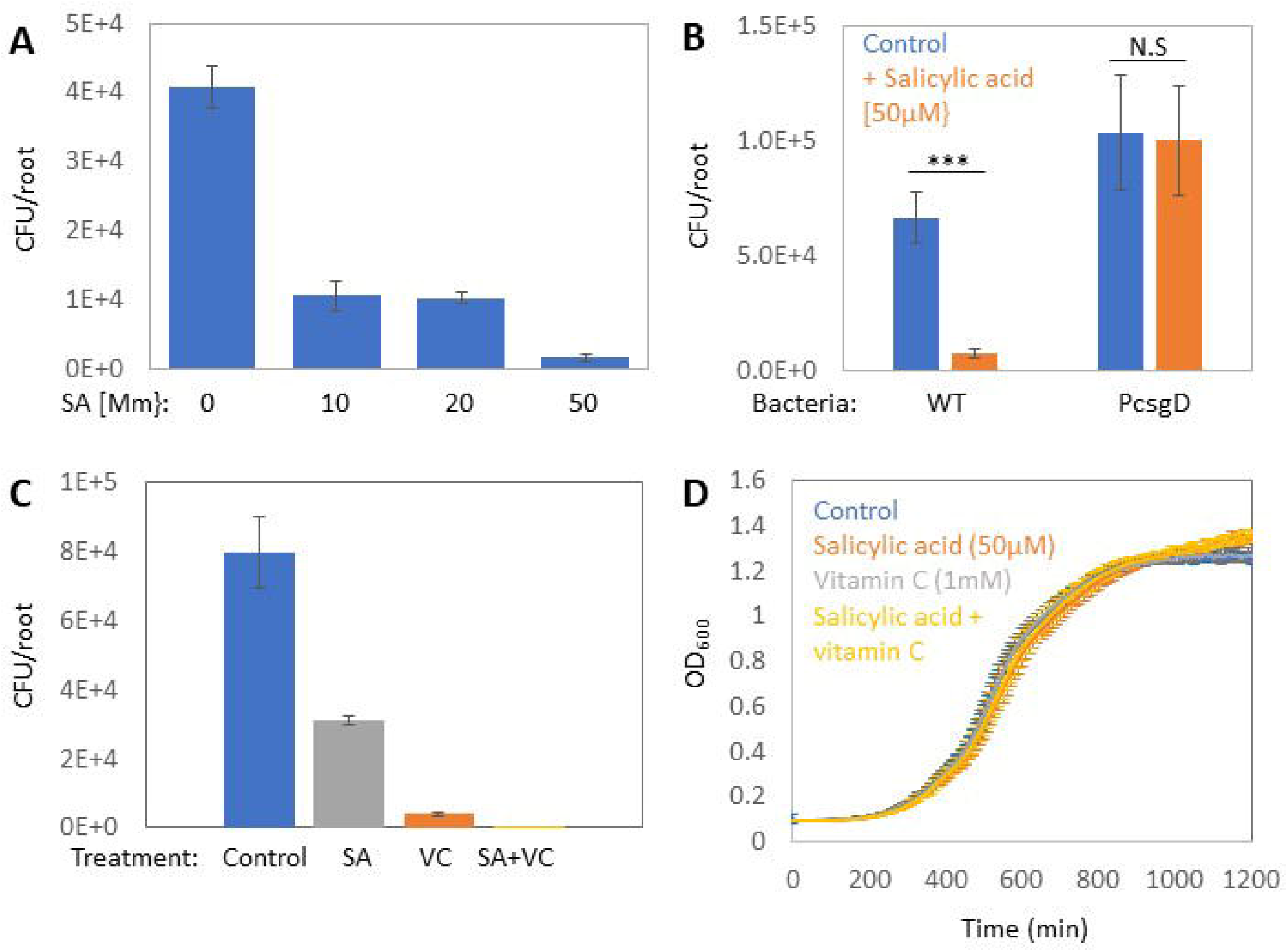
Salicylic acid and vitamin C synergistically inhibit root colonization. **(A)** 7 days old seedlings were inoculated with *Salmonella* on MS agar plates with or without the indicated concentration of salicylic acid, and the number of colonizing bacteria was determined after 48 hrs. Shown are averages and standard deviations (SD) of bacterial colonization from 2 independent experiments with 4 seedlings in each, all the treatments significantly reduced bacterial colonization P<0.005. **(B)** 7 days old seedlings were inoculated with the indicated *Salmonella* strains on MS agar plates with or without salicylic acid 50μM, and the number of colonizing bacteria was determined after 48 hrs. Shown are averages and standard deviations (SD) of bacterial colonization from 3 independent experiments with 4 seedlings in each, ***=P<0.005, N.S= non-significant. **(C)** seedlings were inoculated bacteria on MS agar plates with the indicated chemicals: 50μM salicylic acid, 1mM vitamin C and a combination of both, and the number of colonizing bacteria was determined after 48 hrs. Shown are averages and standard deviations (SD) of bacterial colonization from .independent experiments with 3 seedlings in each. ***=P<0.005. **(D)** Bacteria were grown in LB with the addition of the indicated chemicals and OD_600_ measured. Shown are averages and SD of 5 independent cultures for each treatment.

The genus *Salmonella* harbors more than two thousand different serovars, with varying abilities cause disease in humans. The combination of vitamin C and SA significantly inhibited the colonization of 11 different serovars isolated from human patients we tested (Figure 4A), with inhibition efficiency ranging from 60% (*S. virginia*) to 95%-99% in most other serovars. We next examine if SA and vitamin C can inhibit plant colonization of other human enteric pathogens associated with plant food contamination. Examination root colonized by enteropathogenic *E*.*coli* (EPEC serotype O127:H6) ^29^ and the gram-positive pathogen *Listeria monocytogenes* (strain 10403s) revealed the efficiency of vitamin C and SA and the synergistic effect of their combination for inhibiting these bacteria as well (Figure 4B-4C).

**Figure 4.**
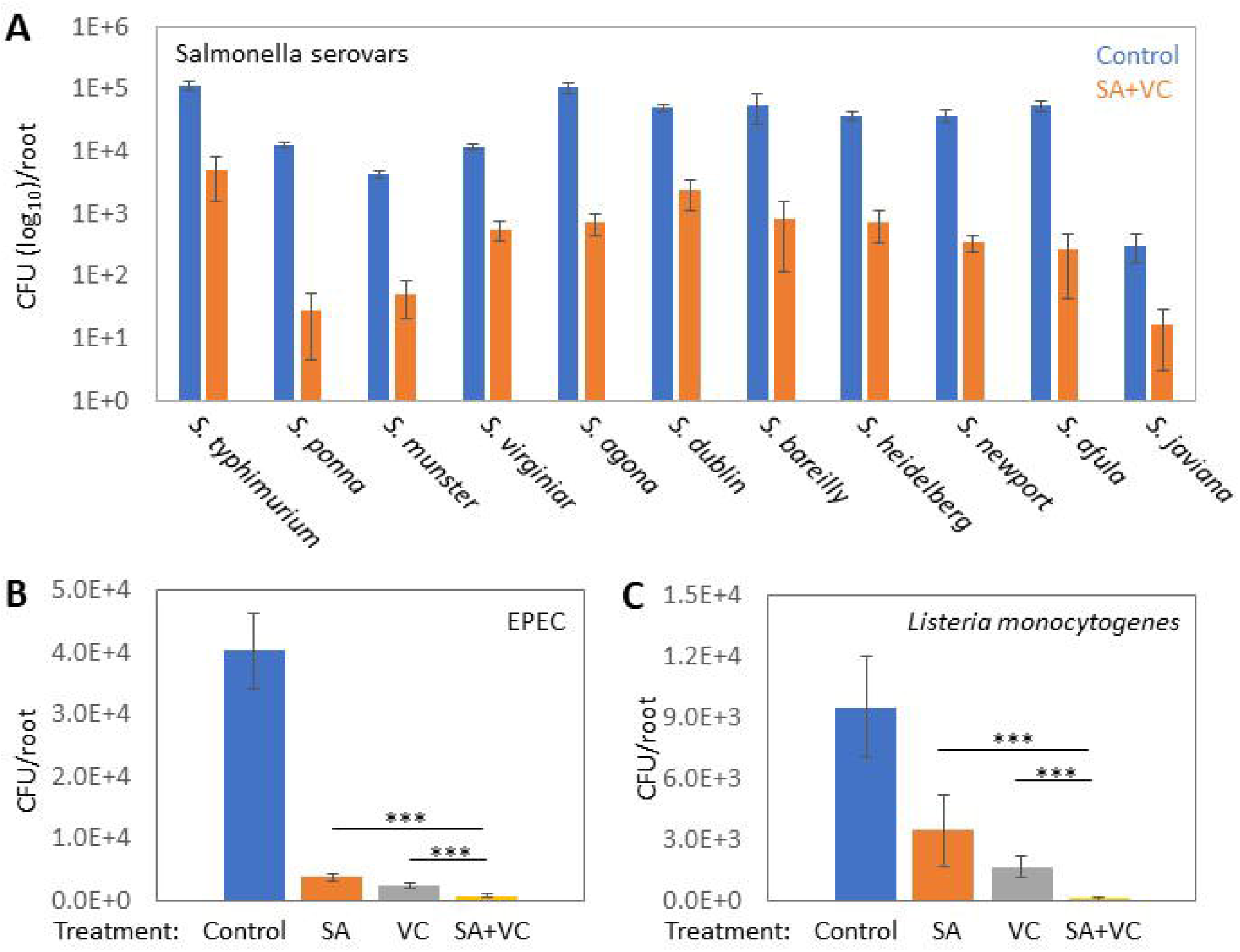
SA and VC affect root colonization by diverse human pathogens. **(A)** 7 days old seedlings were inoculated with the indicated *Salmonella* serovars on MS agar plates with or without a combination of 50μM salicylic acid and 1mM vitamin C. The number of colonizing bacteria was determined after 48 hrs. Shown are averages and standard deviations (SD) of bacterial colonization from at least 4 seedlings for each serovar. For all the serovars, the P<0.005, except for *S. Bareilly* and *S*. Havana, where P<0.05. (**B-C**) Seedlings were inoculated with either enteropathogenic *E. coli* (O127:H6) (B) or *Listeria monocytogenes* (10403s) with the indicated chemicals: 50μM salicylic acid, 1mM vitamin C and a combination of both, and the number of colonizing bacteria determined after 48hrs. Shown are averages and standard deviations (SD) of bacterial colonization from 3 independent experiments with 3 seedlings in each. ***=P<0.005.

Next, we explored the effect of vitamin C and SA on *salmonella* colonization of edible plants. Using sterile alfalfa sprouts and Basil leaves (Ocimum basilicum), we confirmed the effect of vitamin C and SA on *salmonella* contamination of these economically important foods (Figure 5A-5B). Thus, the effect is extended beyond Arabidopsis to edible plants and beyond the root to plant leaves, highlighting its general effect. Notably, neither vitamin C nor SA inhibits leaf colonization alone, but the combination efficiently inhibited colonization (Figure 5B).

**Figure 5.**
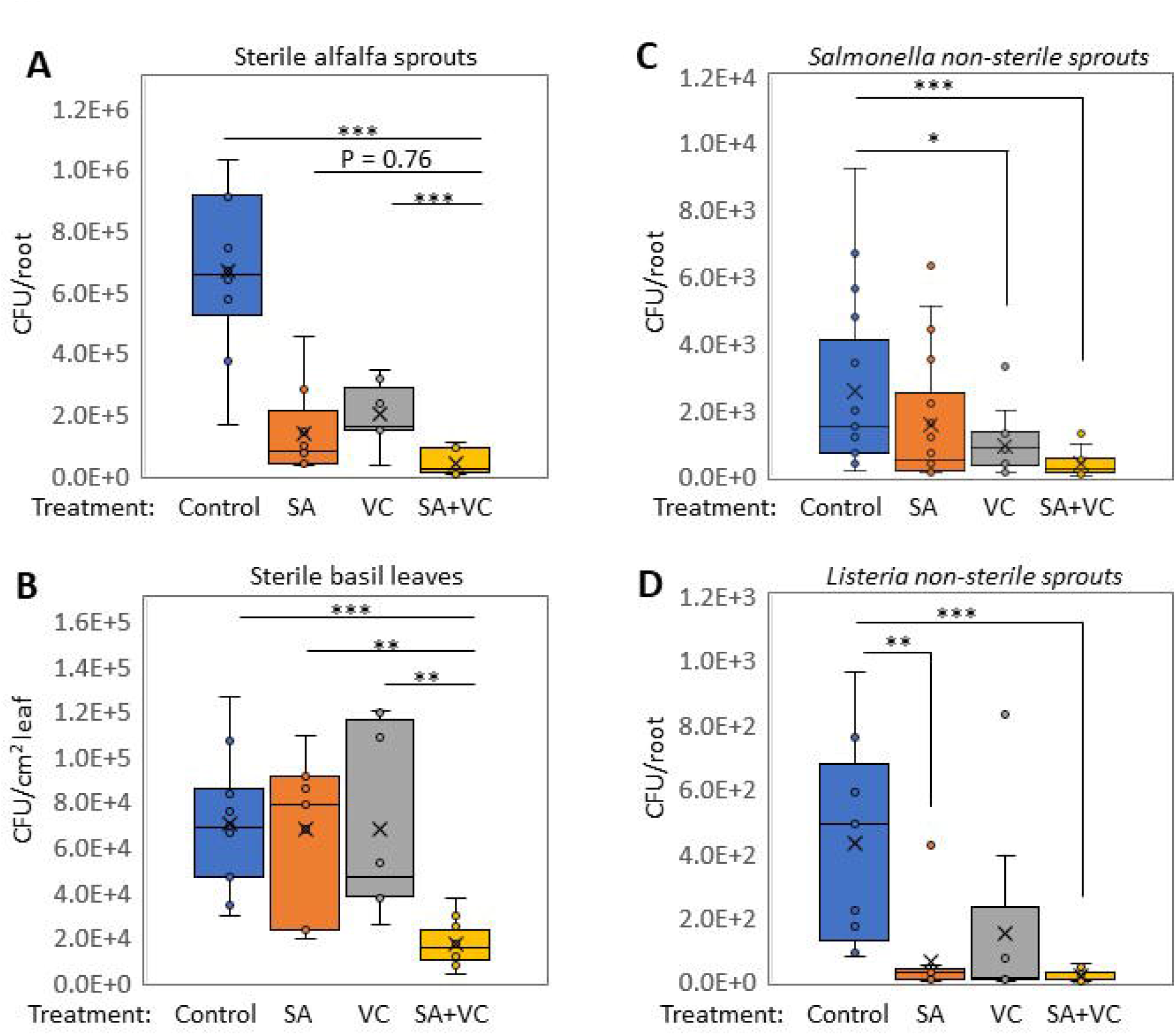
SA and VC affect basil leaves, and alfalfa sprout colonization. **(A)** 5 days old alfalfa seedlings were inoculated bacteria on MS agar plates with the indicated chemicals: 50μM salicylic acid, 1mM vitamin C and a combination of both, and the number of colonizing bacteria was determined after 48 hrs. Shown are averages and standard deviations (SD) of bacterial colonization from 3 independent experiments with 3 seedlings in each. ***=P<0.005. **(B)** Leaves excised from 5 sterile grown 5 weeks old basil and placed on MS agar plates with the indicated chemicals as described in (A) and the number of colonizing bacteria determined after 48hrs. Shown are averages and standard deviations (SD) of bacterial colonization from 3 independent experiments with 3 seedlings in each. **=P<0.01, ***=P<0.005. (**C-D**) nonsterile alfalfa sprouts were inoculated with GFP expressing *Salmonella* (C) and *Listeria* (D) with the indicated chemicals as described in (A), and the number of colonizing bacteria was determined after 48 hrs after plating on semi-selective plates and examination under the microscope. Shown are averages and standard deviations (SD) of bacterial colonization from 3 independent experiments with 3 seedlings in each. *=P<0.05, **=P<0.01, ***=P<0.005.

The experiments so far have been done on sterile plants grown in axenic conditions without resident microbiota. We further examined the effect of vitamin C and SA on Salmonella while colonizing non-sterile alfalfa sprouts. The colonization of nontreated plants by *salmonella* was reduced by two orders of magnitude compared to sterile plants [compare Figure 5A to Figure 5C], suggesting that competition with the resident microbiota interferes with the pathogen’s colonization. Nevertheless, vitamin C and SA further inhibited the colonization (Figure 5C). Similar inhibition was observed for *Listeria monocytogenes* (Figure 5D). Thus, our data suggest that SA and vitamin C or the combination of both is effective way to prevent plant contamination by enteric pathogens.

We finally wondered if other bacteria may exploit the plant immune system to enhance their colonization the same way we found in Salmonella. To examine this possibility, we monitored root colonization by a collection of bacteria from the *Pseudomonadota* phylum isolated from diverse environments. We found isolates that exploit the plant immune system, as suggested by the reduction in colonization on bbc triple mutant roots (Figure S4). Interestingly, all the 3 isolates that seem to exploit the plant immune system were isolated from the leaf niche and may evolve to adapt to the plant environment. However, a larger sample size and a careful comparison are necessary to support such a conclusion. Anyway, our result suggest that other bacteria can also exploit the plant immune system to enhance their colonization.

## Discussion

*Salmonella E*.*coli* and *Listeria* are the three main bacteria causing contamination of fresh produce, leading to enormous health and economic burdens. In order to develop sustainable and environmentally friendly methods to control fresh produce contamination by enteric pathogens, we need a mechanistic understanding of their interaction with plants. Our research revealed that *salmonella* exploits the plant immune system activation, using plant-produced ROS to stimulate biofilm formation and root adhesion. Interestingly, biofilm protects the bacteria from the plant ROS, suggesting a feedback loop where 1. colonization stimulates immune system activation (Figure 1A). 2. The immune system produces ROS to inhibit the bacteria. 3. Bacteria can utilize ROS to enhance biofilm formation (Figure 2B) 4. Biofilm protects bacteria from the immune system (Figure 2C and Figure 3A) and further enhances colonization [see model in (Figure 6)].

**Figure 6.**
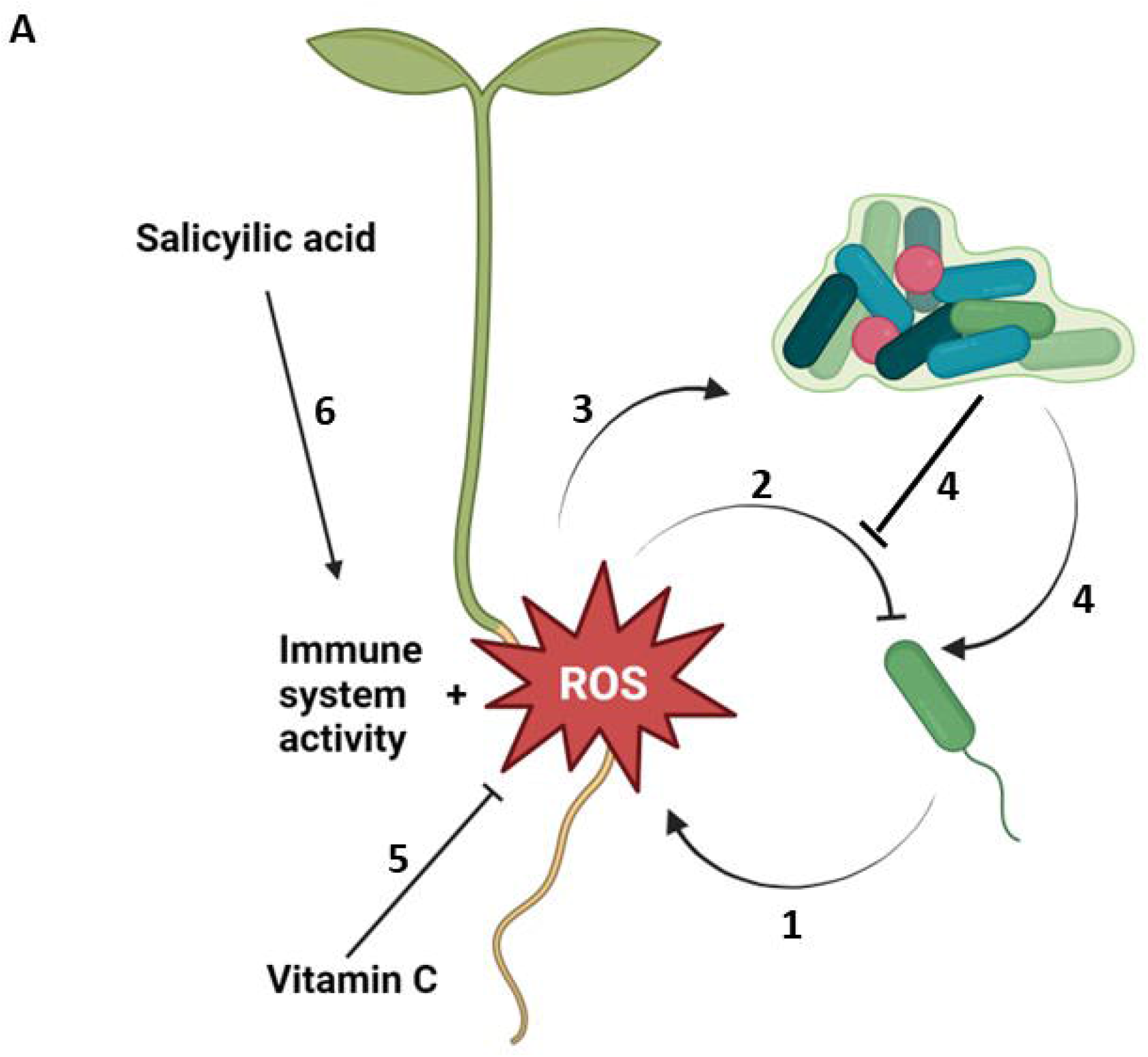
A model describing how *Salmonella* exploits the plant immune system and how we can outsmart it. **(A)** The plant recognizes the *Salmonella* and activates an immune response and ROS production (1) to inhibit bacterial colonization (2). *Salmonella* exploits the ROS for enhancing biofilm formation and promoting root colonization (3). The biofilm formation further protects the bacteria from the immune system activity and ROS toxicity (4). Reducing ROS by antioxidants like VC inhibits colonization by interfering with biofilm formation (5) and, in turn, makes the bacteria vulnerable to the strong immune system activated by SA (6). Thus, VC and SA exhibit synergistic activity in preventing plant contamination by *Salmonella* and other pathogens.

Previous research claimed that *Salmonella* can infiltrate the plant endophytic compartment and exhibit plant pathogenicity phenotype^11,25^. Our results suggest that pathogenic phenotypes only manifested at high bacterial concentrations, while low concentration exhibit beneficial effects (Figure S2A). The difference in the number of inoculated bacteria may also explain why we failed to observe endophytic infiltration in our system. Remarkably, it was recently demonstrated that non-pathogenic bacteria may elicit pathogenic like effects in Arabidopsis leaves when inoculated at high concentrations^30^. Our data suggest that Salmonella face extensive competition from the resident plant microbiota (compare figure 5A to figure 5D) and the number of *Salmonella* is reduced to ∼100 bacteria per sprout when colonized non-sterile sprouts suggesting that 10^5^ CFU is an unlikely scenario in the real world.

Work on *Salmonella* infection in human and mice also found indications that salmonella might exploit the ROS produced by the immune system^31^. However, such a phenomenon has not been described in plant. We have previously demonstrated that immune system activation enhances *Bacillus veleznesis* root adhesion^23^. The current work further highlights the complex interaction between bacteria and the immune system, beyond merely bacterial killing or growth inhibition.

We further found that vitamin C and salicylic acid synergistically inhibited salmonella colonization, we hypothesize that vitamin C inhibit biofilm formation exposing the bacteria to the immune system activity stimulated by SA like proteolytic enzymes, antimicrobial molecules etc ^32^. Our work demonstrates how mechanistic understanding of the complex interaction between bacterial colonization and plant immunity opens a new avenue for dealing with food contamination. We discovered that safe-to-use antioxidants like vitamin C can inhibit plant colonization by enteric pathogens and synergize with immune system activation by salicylic acid. The results from the root of the model plant arabidopsis were also applied to basil leaves and alfalfa sprouts, even under non-sterile conditions in the presence of resident microbiota. Vitamin C and low doses of salicylic acid are safe for human consumption and have other health benefits on their own^33,34^. Further exploring the utilization of this combination on fresh produce in real world settings is a promising avenue for sustainable food safety.

## Acknowledgments

We thank Prof. Edward Miao (Duke), Prof Sheng Yang He (Duke), Prof Ferric Fang (WU), Prof. Niko Geldner (UNIL), Prof Anat Herskovits (TAU), Prof Ute Romling (KI) for bacterial strains and Arabidopsis lines. The research was funded by the Israeli Department of Agriculture Grant code: 21-01-0075.

In memory of Prof. Philip Benfey a great mentor and person.

## Material and methods

### Plant and bacterial strains

are listed in Table S1

#### Plant growth

Arabidopsis seedlings and sterile alfalfa seedlings were grown on 0.5 MS media containing 1.1 gr Murashige and Skoog basal salts (in 500 ml ddH2O), 1% sucrose, 1% agar and 5 ml (in 500 ml ddH2O) MES (50 gr/l, pH=5.8 with NaOH). Arabidopsis seeds were stratified for 2 d in a 4C dark room and grown vertically for 7 days under long-day light conditions. Sterile alfalfa seedlings were grown vertically for 5 days under long-day light conditions. Non-sterile sprouts were bought in the local market. Basil plants were grown in a sterile box on 0.5 MS media for 5 weeks under long-day light conditions and then leaves were excised for the bacterial inoculation experiments.

#### Monitoring bacterial growth and plant colonization

Bacterial inoculation and CFU measurement was done as described previously ^35^. Briefly, fresh bacterial colonies were grown at 250 RPM, shaking overnight in LB media at 23°. The next day, the culture was diluted 1:500 in fresh LB and grown at 37° for OD_600_=1. We washed the bacteria in 10mM MgCl_2_ buffer and then diluted the culture 1:100 in 10mM MgCl_2_. 2μL of the diluted culture was gently applied on the tip of the root/stem (alfalfa)/cut leaf surface (basil) and placed on an ½ MS agar plate with no sucrose. For figure S1A (2) bacteria were added to the MS media before pouring the plates to the concentration denoted. The square plates were kept in a vertical position during the incubation time at 23° under long-day light conditions (16 h light/8 h darkness) in a plant growth chamber. For bacterial CFU counting and microscopy, plants were incubated with bacteria for 48 hrs. Then, the inoculated plant roots were cut and washed three times in sterile water. The seedlings were transferred to a tube with 1 ml of MgCl_2_ and vortexed vigorously for 20 seconds, then serial dilutions were plated on LB plates for CFU counting or observed under confocal microscope Zeiss LSM980. The whole root and stem were taken from Arabidopsis and alfalfa, while a circle cut of 5mm in diameter was taken from the basil leaves. For colonization of non-sterile sprouts, we used GFP *Salmonella* and GFP *Listeria* and plated the samples on Kanamycin plates for the Salmonella and Listeria selective enrichment supplement containing plates (Sigma) for the Listeria. The identity of the bacteria was validated by monitoring the GFP expression under a binocular (Leica). When indicated, the following chemicals were added to the MS media: salicylic acid, KI, Diphenyl iodonium (DPI) all the chemicals purchased from Sigma Aldrich. The pH of the MS was titrated back to normal after the addition of salicylic acid and ascorbic acid with NaOH. For measuring endophytic colonization seedlings were grown with *Salmonella* as described above, after 48 hrs the seedlings were surface sterilized for 5 minutes in 10% hypochlorite + 1% Tween 20. Then the roots were washed 3 times in sterilized water, the root were crushed in 10mM MgCl_2_ and plated on LB agar plates.

Bacterial growth in the presence SA and vitamin C was measured in 48 well plates. Overnight cultures were diluted into fresh media with or without the chemicals, and OD_600_ was measured every 10 minutes for 12 hours in Tecan spark 10M.

#### Biofilm formation assay

Quantification of biofilm formation was done as described previously ^36^. Briefly, the overnight culture of bacteria was diluted into LB with no NaCl in a multi-well plate. The plates were incubated at 23° for 5 days without shaking. After 5 days the media was poured, and a solution of 0.1% crystal violet was added and incubated for 10 min. Next, the wells were washed 3 times with tape water and dried on the bench. A solution of 33% acetic acid was added to dissolve the crystal violet bound to the wall, and the OD_570_ was measured.

#### Pseudomonas isolation

We isolated *pseudomonases* from the local area (Kiryat Shmona, north of Israel) from 3 different niches: Local soil, local stream, and mint leaves. The soil was vigorously shaken in LB. The leaf was ground in LB. The samples were plated on Pseudomonas isolation agar (Sigma) for 48 hours at 30°. Colonies were re-streaked on Pseudomonas isolation agar to get pure colonies, and the bacterial identity was determined by 16S sequencing (Table S1).

#### Data analysis

Root length was manually measured in imageJ. Statistical analysis (student T-tset) was done in Excel. Data representation in graphs was done in Excel.

## Notes

### Competing Interest Statement

The authors have declared no competing interest.

### Summary of Updates

New experiments conducted to improve the data

## References

1 Benaissa, A. Rhizosphere: Role of bacteria to manage plant diseases and sustainable agriculture-A review. J Basic Microbiol (2023). 10.1002/jobm.202300361

2 Zdolec, N., Lorenzo, J. M. & Ray, R. C. Use of Microbes for Improving Food Safety and Quality. Biomed Res Int 2018, 3902698 (2018). 10.1155/2018/3902698

3 https://www.who.int/news-room/fact-sheets/detail/food-safety.

4 Kowalska, B. Fresh vegetables and fruit as a source of Salmonella bacteria. Ann Agric Environ Med 30, 9–14 (2023). 10.26444/aaem/156765

5 Truong, H. N. et al. Plants as a realized niche for Listeria monocytogenes. Microbiologyopen 10, e1255 (2021). 10.1002/mbo3.1255

6 George, A. S. & Brandl, M. T. Plant Bioactive Compounds as an Intrinsic and Sustainable Tool to Enhance the Microbial Safety of Crops. Microorganisms 9 (2021). 10.3390/microorganisms9122485

7 Lim, J. A., Lee, D. H. & Heu, S. The interaction of human enteric pathogens with plants. Plant Pathol J 30, 109–116 (2014). 10.5423/PPJ.RW.04.2014.0036

8 Melotto, M., Panchal, S. & Roy, D. Plant innate immunity against human bacterial pathogens. Front Microbiol 5, 411 (2014). 10.3389/fmicb.2014.00411

9 Garcia, A. V. et al. Salmonella enterica flagellin is recognized via FLS2 and activates PAMP-triggered immunity in Arabidopsis thaliana. Mol Plant 7, 657–674 (2014). 10.1093/mp/sst145

10 Roy, D., Panchal, S., Rosa, B. A. & Melotto, M. Escherichia coli O157:H7 induces stronger plant immunity than Salmonella enterica Typhimurium SL1344. Phytopathology 103, 326–332 (2013). 10.1094/PHYTO-09-12-0230-FI

11 Schikora, A., Carreri, A., Charpentier, E. & Hirt, H. The dark side of the salad: Salmonella typhimurium overcomes the innate immune response of Arabidopsis thaliana and shows an endopathogenic lifestyle. PLoS One 3, e2279 (2008). 10.1371/journal.pone.0002279

12 Jones, J. D. & Dangl, J. L. The plant immune system. Nature 444, 323–329 (2006). 10.1038/nature05286

13 Xin, X. F., Kvitko, B. & He, S. Y. Pseudomonas syringae: what it takes to be a pathogen. Nat Rev Microbiol 16, 316–328 (2018). 10.1038/nrmicro.2018.17

14 Fitzpatrick, C. R. et al. The Plant Microbiome: From Ecology to Reductionism and Beyond. Annu Rev Microbiol 74, 81–100 (2020). 10.1146/annurev-micro-022620-014327

15 Yaron, S. & Romling, U. Biofilm formation by enteric pathogens and its role in plant colonization and persistence. Microb Biotechnol 7, 496–516 (2014). 10.1111/1751-7915.12186

16 Flemming, H. C. et al. The biofilm matrix: multitasking in a shared space. Nat Rev Microbiol 21, 70–86 (2023). 10.1038/s41579-022-00791-0

17 Bano, S. et al. Biofilms as Battlefield Armor for Bacteria against Antibiotics: Challenges and Combating Strategies. Microorganisms 11 (2023). 10.3390/microorganisms11102595

18 Morris, C. E. & Monier, J. M. The ecological significance of biofilm formation by plant-associated bacteria. Annu Rev Phytopathol 41, 429–453 (2003). 10.1146/annurev.phyto.41.022103.134521

19 Zhou, F. et al. Co-incidence of Damage and Microbial Patterns Controls Localized Immune Responses in Roots. Cell 180, 440–453 e418 (2020). 10.1016/j.cell.2020.01.013

20 Millet, Y. A. et al. Innate immune responses activated in Arabidopsis roots by microbe-associated molecular patterns. Plant Cell 22, 973–990 (2010). 10.1105/tpc.109.069658

21 Yuan, M. et al. Pattern-recognition receptors are required for NLR-mediated plant immunity. Nature 592, 105–109 (2021). 10.1038/s41586-021-03316-6

22 Pfeilmeier, S. et al. The plant NADPH oxidase RBOHD is required for microbiota homeostasis in leaves. Nat Microbiol 6, 852–864 (2021). 10.1038/s41564-021-00929-5

23 Tzipilevich, E., Russ, D., Dangl, J. L. & Benfey, P. N. Plant immune system activation is necessary for efficient root colonization by auxin-secreting beneficial bacteria. Cell Host Microbe 29, 1507–1520 e1504 (2021). 10.1016/j.chom.2021.09.005

24 Torres, M. A., Jones, J. D. & Dangl, J. L. Pathogen-induced, NADPH oxidase-derived reactive oxygen intermediates suppress spread of cell death in Arabidopsis thaliana. Nat Genet 37, 1130–1134 (2005). 10.1038/ng1639

25 Iniguez, A. L. et al. Regulation of enteric endophytic bacterial colonization by plant defenses. Mol Plant Microbe Interact 18, 169–178 (2005). 10.1094/MPMI-18-0169

26 Newman, S. L., Will, W. R., Libby, S. J. & Fang, F. C. The curli regulator CsgD mediates stationary phase counter-silencing of csgBA in Salmonella Typhimurium. Mol Microbiol 108, 101–114 (2018). 10.1111/mmi.13919

27 Ahmad, I. et al. Complex c-di-GMP signaling networks mediate transition between virulence properties and biofilm formation in Salmonella enterica serovar Typhimurium. PLoS One 6, e28351 (2011). 10.1371/journal.pone.0028351

28 Grantcharova, N., Peters, V., Monteiro, C., Zakikhany, K. & Romling, U. Bistable expression of CsgD in biofilm development of Salmonella enterica serovar typhimurium. J Bacteriol 192, 456–466 (2010). 10.1128/JB.01826-08

29 Levine, M. M. et al. Escherichia coli strains that cause diarrhoea but do not produce heat-labile or heat-stable enterotoxins and are non-invasive. Lancet 1, 1119–1122 (1978). 10.1016/s0140-6736(78)90299-4

30 Miebach, M., Faivre, L., Schubert, D., Jameson, P. & Remus-Emsermann, M. Nonpathogenic leaf-colonizing bacteria elicit pathogen-like responses in a colonization density-dependent manner. Plant Environ Interact 5, e10137 (2024). 10.1002/pei3.10137

31 Rhen, M. Salmonella and Reactive Oxygen Species: A Love-Hate Relationship. J Innate Immun 11, 216–226 (2019). 10.1159/000496370

32 Yalpani, N., Silverman, P., Wilson, T. M., Kleier, D. A. & Raskin, I. Salicylic acid is a systemic signal and an inducer of pathogenesis-related proteins in virus-infected tobacco. Plant Cell 3, 809–818 (1991). 10.1105/tpc.3.8.809

33 Suliburska, J. & Cholik, R. S. Risks and benefits of salicylates in food: a narrative review. Nutr Rev (2023). 10.1093/nutrit/nuad136

34 Mousavi, S., Bereswill, S. & Heimesaat, M. M. Immunomodulatory and Antimicrobial Effects of Vitamin C. Eur J Microbiol Immunol (Bp) 9, 73–79 (2019). 10.1556/1886.2019.00016

35 Tzipilevich, E. & Benfey, P. N. Phage-Resistant Bacteria Reveal a Role for Potassium in Root Colonization. mBio 12, e0140321 (2021). 10.1128/mBio.01403-21

36 Trampari, E. et al. Exposure of Salmonella biofilms to antibiotic concentrations rapidly selects resistance with collateral tradeoffs. NPJ Biofilms Microbiomes 7, 3 (2021). 10.1038/s41522-020-00178-0

