## Supplementary figures and images for "Salmonella exploits the reactive oxygen species generated by the plant immune system to enhance colonization"

### Figure S1

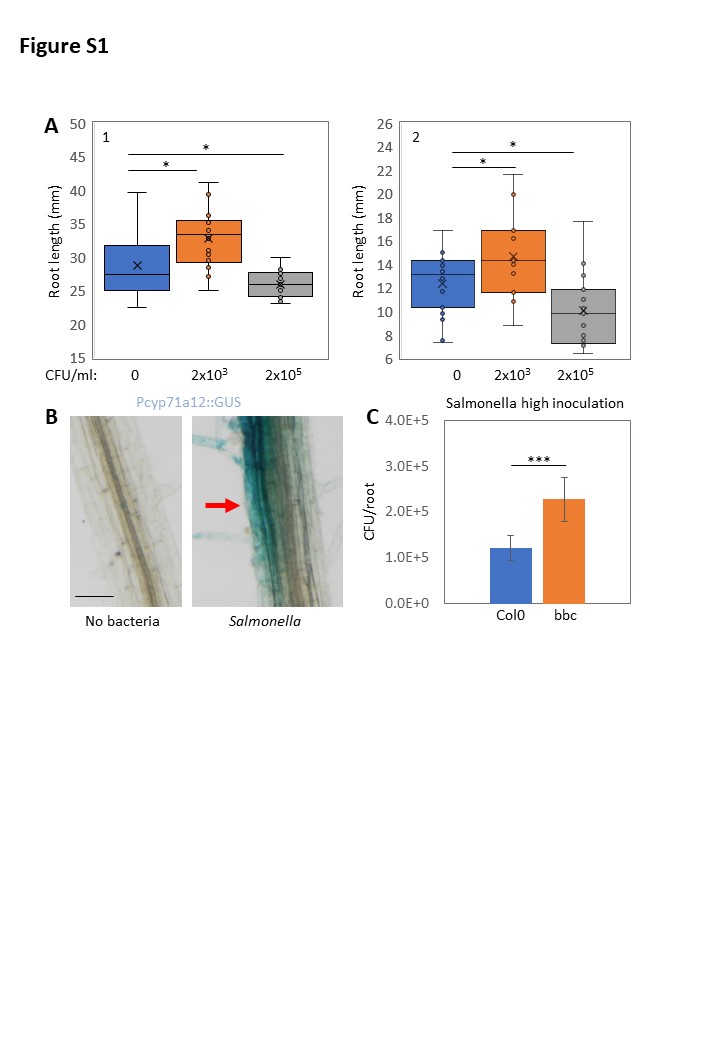

### Figure S2

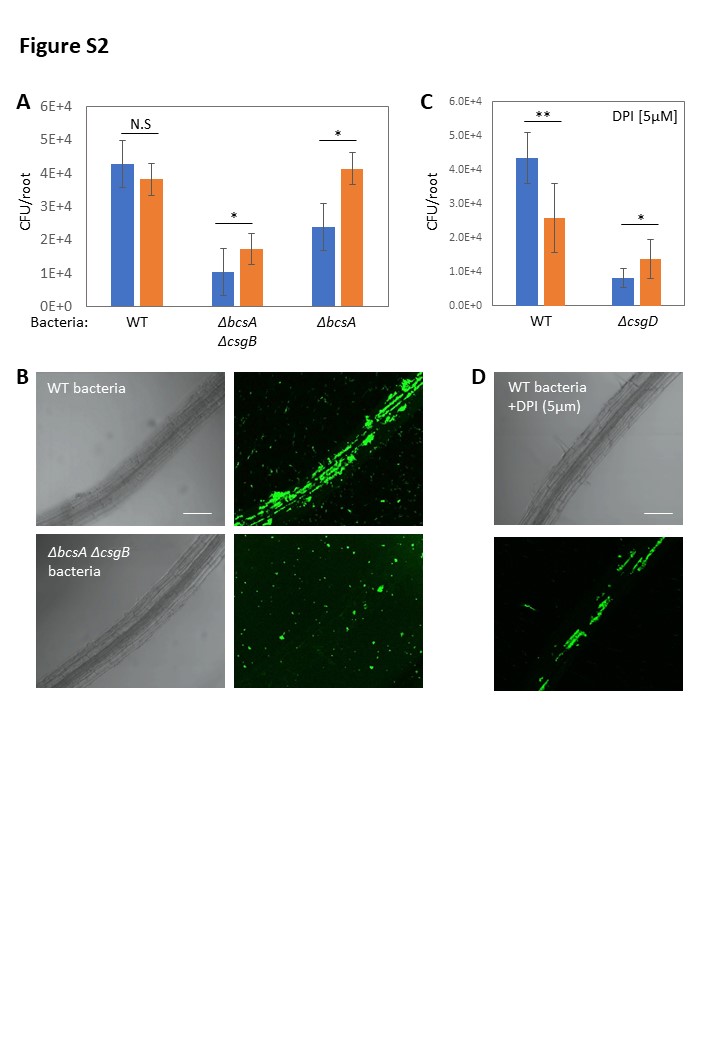

### Figure S3

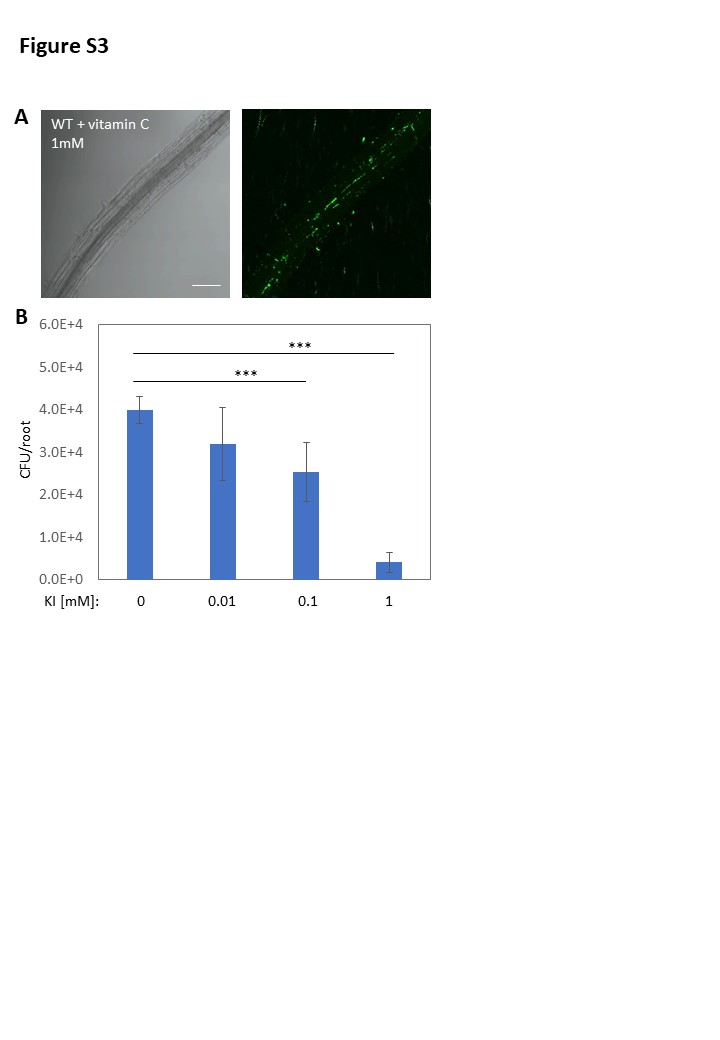

### Figure S4

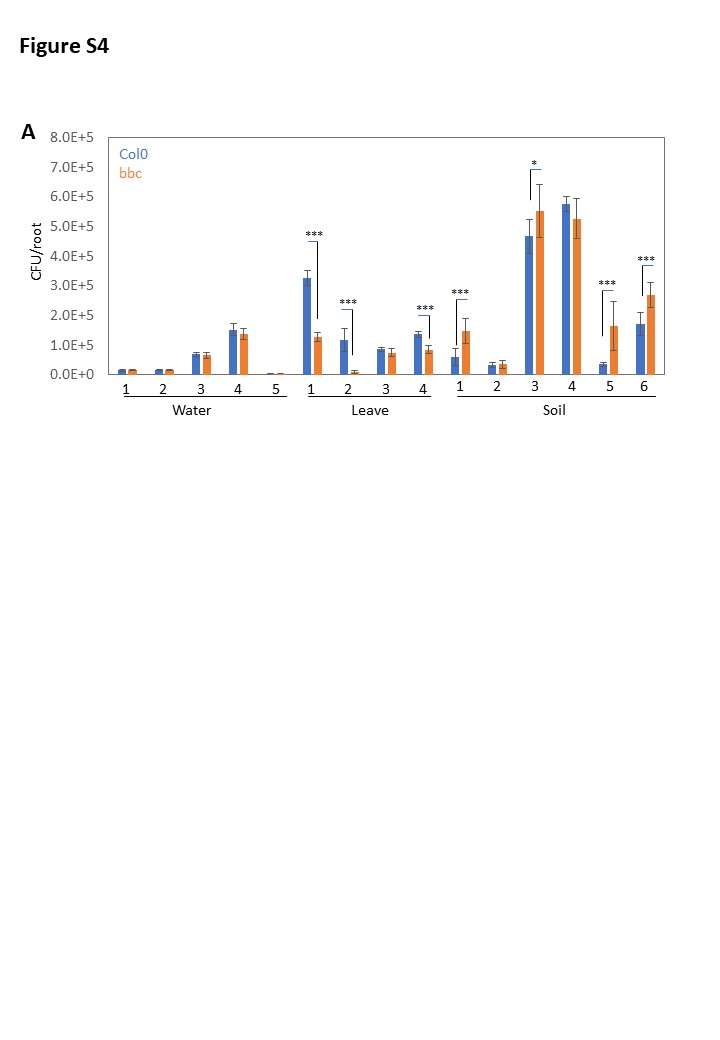
