## Supplementary figure legends for "Salmonella exploits the reactive oxygen species generated by the plant immune system to enhance colonization"

**Figure S1. Low number of Salmonella exhibit plant beneficial effect and activate immune system**

**(A)** 1. 7 days old WT (col0) seedlings were inoculated with Salmonella with the indicated concentration and root length measured after 48 hours. 2. Sterilized Arabidopsis seeds were sown in the presence of the indicated bacterial concentrations and root length measured. Shown are whisker plots of root length, * = P < 0.05.

**(B)** 7 days old Arabidopsis seedlings (Pcyp71a12::GUS) were inoculated with bacteria on MS agar plates and GUS staining was monitored after 18 hrs . Shown are representative images out of 4 roots from each treatment, scale bar = 25μm.

**(C)** Seedlings of the indicated genotypes were inoculated with *Salmonella* ($10^8$ CFU/ml), and the number of colonizing bacteria was determined after 48 hrs. Shown are averages and standard deviations (SD) of bacterial colonization from at least 2 independent experiments with 4 seedlings in each. *** = P < 0.005.

**Figure S2. Plant immunity reduced the colonization of biofilm mutant bacteria**

**(A)** Seedlings of either WT (Col0) or bbc mutants were inoculated with the indicated *Salmonella* strains UMRI (rdar morphotype[27]) used as control. The number of colonizing bacteria was determined after 48 hrs. Shown are averages and standard deviations (SD) of bacterial colonization from at least 2 independent experiments with 4 seedlings in each. * = P < 0.05, N.S = non-significant.

**(B)** WT (Upper panels) and *ΔcsgB ΔbcsA* double mutant *Salmonella* (lower panels) constitutively expressing GFP were inoculated into seedlings of the indicated genotypes for 48 hrs. Root colonization was monitored by confocal microscopy. Shown are representative maximal projection x20 fluorescent images and the complementary brightfield images of the same roots, out of 4 roots from each treatment. Scale bar = 50μm.

**(C)** WT (Col0) seedlings were inoculated with the indicated *Salmonella* strains in the presence or absence of 5μM DPI (diphenyliodonium) an inhibitor of NADPH oxidases. The number of colonizing bacteria was determined after 48 hrs. Shown are averages and standard deviations (SD) of bacterial colonization from at least 2 independent experiments with 4 seedlings in each. ** = P<0.01, * = P < 0.05.

**(D)** WT *Salmonella* constitutively expressing GFP were inoculated into seedlings treated with 5mM DPI for 48 hrs. Root colonization was monitored by confocal microscopy. Shown is a representative maximal projection x20 fluorescent image (lower panel) and the complementary brightfield image (upper panel) of the same root, out of 4 roots with the same treatment. Scale bar = 50μm.

**Figure S3. The antioxidants vitamin C and KI significantly reduced *Salmonella* colonization**

**(A)** WT *Salmonella* constitutively expressing GFP were inoculated into seedlings treated with 1mM vitamin C for 48 hrs. Root colonization was monitored by confocal microscopy. Shown is a representative maximal projection x20 fluorescent image (lower panel) and the complementary brightfield image (upper panel) of the same root, out of 4 roots with the same treatment. Scale bar = 50μm.

**(B)** 7 days old seedlings were inoculated with *Salmonella* on MS agar plates with or without the indicated concentration of potassium iodide (KI), and the number of colonizing bacteria was determined after 48 hrs. Shown are averages and standard deviations (SD) of bacterial colonization from 2 independent experiments with 4 seedlings in each. *** = P<0.005.

**Figure S4. Bacteria from the *pseudomonas* genus can exploit the plant immune system to enhance colonization.**

**(A)** 7 days old Col0 and bbc seedlings were inoculated with bacteria from the *Pseudomonadota* phylum isolated from 3 different habitats (see Table S1 for the bacterial identity as determined by 16s sequencing). The number of colonizing bacteria was determined after 48 hrs. Shown are averages and standard deviations (SD) of bacterial colonization from at least 4 seedlings for each bacterium.
