## Supplementary material for "Salmonella exploits the reactive oxygen species generated by the plant immune system to enhance colonization": Table S1

**Table S1. Bacterial and plant strains**

| **Bacterial strains** | | |
|---|---|---|
| **Strain Name** | **Description** | **Reference and Comments** |
| **Salmonella strains** | | |
| *Salmonella enterica 14028s* | *Salmonella enterica subsp. enterica ATCC 14028s* | Kindly provided by Prof. Edward Miao Duke university |
| *Salmonella enterica 14028s ΔflgB* | *flgB::*Tn10 [37] | Kindly provided by Prof. Edward Miao Duke university |
| *Salmonella enterica 14028s* GFP | Wild type *S. enterica* 14028s strain with constitutive GFP expression | Kindly provided by Prof. Edward Miao Duke university |
| *Salmonella enterica 14028s ΔcsgD* | SLN112 [26] | Kindly provided by Prof. Ferric Fang Washington university |
| *Salmonella enterica 14028s* pCsgD | SLN135 [26] | Kindly provided by Prof. Ferric Fang Washington university |
| *Salmonella enterica 14028s* RDAR morphotype | UMR1 [28] | Kindly provided by Ute Romling Karolinska institute |
| *Salmonella enterica 14028s* RDAR morphotype *ΔbcsA* | MAE150 [28] | Kindly provided by Ute Romling Karolinska institute |

| | | |
|---|---|---|
| *Salmonella enterica 14028s* RDAR morphotype *ΔbcsA, ΔcsgB* | MAE190 [28] | Kindly provided by Ute Romling Karolinska institute |
| **Other bacterial strains** | | |
| E2348/69 (*wt*) | EPEC-*wt* isolate, serotype O127:H6, NA | J. Kaper, [29] |
| *Listeria monocytogenes 10403s* | WT isolate | Kindly provided by Porf. Anat Herskovits Tel Aviv university |
| GFP *Listeria monocytogenes 10403s* | Wild type L. monocytogenes 10403s strain with constitutive GFP expression | Kindly provided by Porf. Anat Herskovits Tel Aviv university |
| **Isolated pseudomonases** | | |
| *Aeromonas sp. strain VNR183* | Water isolate 1 | Determined by 16s RNA sequencing |
| *Stenotrophomonas maltophilia strain A1-1* | Water isolate 2 | Determined by 16s RNA sequencing |

| | | |
|---|---|---|
| *Pseudomonas alcaliphila strain SS NBRI 36* | Water isolate 3 | Determined by 16s RNA sequencing |
| Pseudomonas punonensis strain JB11 | Water isolate 4 | Determined by 16s RNA sequencing |
| *Stenotrophomonas rhizophila strain S242* | Water isolate 5 | Determined by 16s RNA sequencing |
| *Pseudomonas wadenswilerensis* strain RD2 | Leaf isolate 1 | Determined by 16s RNA sequencing |
| *Pseudomonas sp. JNR-02* | Leaf isolate 2 | Determined by 16s RNA sequencing |
| *Stenotrophomonas sp. strain NJ1024* | Leaf isolate 3 | Determined by 16s RNA sequencing |
| *Pseudomonas putida strain PBS6* | Leaf isolate 4 | Determined by 16s RNA sequencing |
| *Pseudomonas alkylphenolica strain 48A-P9* | Soil isolate 1 | Determined by 16s RNA sequencing |
| *Pseudomonas sp. strain VNP47* | Soil isolate 2 | Determined by 16s RNA sequencing |

| | | |
|---|---|---|
| *Pseudomonas sp. strain huaiR14* | Soil isolate 3 | Determined by 16s RNA sequencing |
| *Pseudomonas donghuensis strain Tn-1* | Soil isolate 4 | Determined by 16s RNA sequencing |
| *Pseudomonas sp. EHE_1_3* | Soil isolate 5 | Determined by 16s RNA sequencing |
| *Pseudomonas fulva strain XL1* | Soil isolate 6 | Determined by 16s RNA sequencing |
| **Plants** | | |
| **Arabidopsis** | | |
| Col0 | WT Arabidopsis line | Lab stock |
| *fls2* | [38] | Lab stock |
| *rbohd: rbohf* | [24] | ABRC |
| bbc | *bak1: bkk1: cerk1* [21] | Kindly provided by Prof. Sheng Yang He Duke university |
| pFrK1::3xmVenus | [19] | Kindly provided by Prof. Niko Geldner university of Lausanne |
| **Other plants** | | |
| Alflafa | *Medicago sativa* | Lab stock |
| Basil | *Ocimum basilicum* | Lab stock |
